# Antibody Reveals Conformational Latch Controlling Herpesvirus Proteases

**DOI:** 10.1101/2025.10.17.683123

**Authors:** Marcell Zimanyi, Kaitlin R. Hulce, Markus-Frederik Bohn, Jordan Norman, Peter J. Rohweder, Yifan Cheng, Charles S. Craik

## Abstract

Human herpesviruses (HHVs) are widespread pathogens that cause severe disease. Their replication depends on the HHV protease (HHV Pr), an enzyme essential for capsid maturation. Because HHV Pr must dimerize to become catalytically active, disrupting dimer formation is a promising strategy for antiviral therapeutic development. We isolated a conformationally selective antibody inhibitor, Fab5, from a fully human naïve Fab-phage library that recognizes monomeric human cytomegalovirus protease (HCMV Pr). A 2.6 Å cryoelectron microscopy (cryo-EM) structure of the Fab5-HCMV Pr complex revealed that Fab5 binds a flexible loop distal from the active site and dimer interface which we call the latch loop. In HCMV Pr dimers, this loop secures the C-terminal tail to the protein core. Structure-guided mutagenesis confirmed that the latch loop is essential for HCMV Pr dimerization and activity. This loop is structurally conserved across all HHV Prs, and we show its functional role in Kaposi’s Sarcoma-associated herpesvirus (KSHV) Pr as well. The latch loop plays a mechanistic role in the conformational transition required for HHV Pr activity, and it forms a cryptic site that presents a new avenue for future allosteric inhibitor development.

## Introduction

Herpesviruses infect over 90% of the human population and establish lifelong latent infections with periodic reactivation^1,2^. Human cytomegalovirus (HCMV) is particularly problematic, causing devasting disease in immunocompromised patients and being the leading global cause of non-genetic congenital malformations, such as deafness and blindness^3,4^. Current treatments suffer from poor efficacy and toxicity, while emergence of treatment-resistant viruses underscore the urgent need for novel therapeutic approaches^5,6^.

Each human herpesvirus (HHV) expresses a serine protease that is essential for viral replication^7,8^. The HHV protease (HHV Pr) activates within the procapsid to cleave scaffolding proteins, leading to capsid maturation (Supplementary Fig. 1). We and others have validated HHV Pr as a therapeutic target by demonstrating that small molecule inhibitors disrupt viral infectivity^9–12^. However, no HHV Pr-targeting compound has advanced through clinical trials. Inhibitor development efforts to date have focused on the active site, which is inherently challenging because it is polar, dynamic and shallow^9^. An orthogonal approach to HHV Pr inhibition is to disrupt the conformational changes required for its activation^5,11^.

All HHV Prs are structurally and functionally conserved and must form homodimers to achieve catalytic activity^13,14^. The micromolar-range dimerization affinities of HHV Prs provide spatial control of proteolysis during viral replication, allowing for dimerization only within the capsid at high local concentration^15^. As a monomer, the HHV Pr C-terminal region is disordered^16–18^. During dimerization, the C-terminus folds into two major helices, one that forms the dimer interface (helix α5) and another which packs beneath the active site (helix α6). This disorder-to-order transition stabilizes the oxyanion hole loop (OHL) which, together with the Ser-His-His catalytic triad, forms an independent active site on each monomer (Fig. 1a)^13^.

**Fig. 1:**
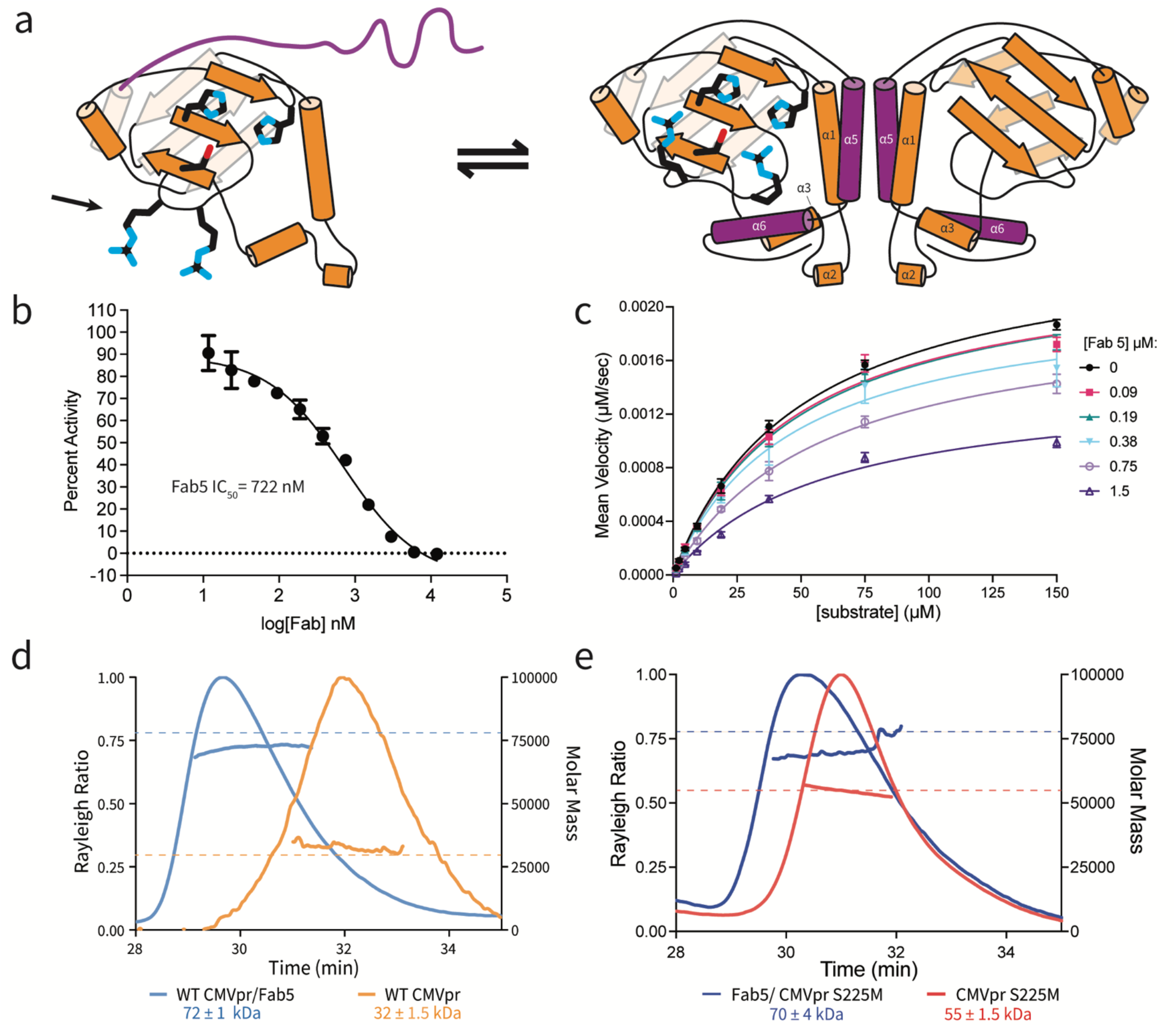
Fab5 is a conformationally selective HCMV Pr inhibitor vesicles. (a) Topology map of HCMV Pr monomer and dimer. In the monomeric state, the C-terminal region (purple) remains disordered, and the oxyanion hole loop (OHL, indicated by arrow) is uncoordinated. Upon dimerization, the C-terminal region adopts a defined structure consisting of helices α5 and α6, with an independent active site on each monomer. (b) Fab5 inhibits HCMV Pr activity with an IC50 of 722 nM. (c) Steady-state kinetics of HCMV Pr performed with increasing concentrations of Fab5. The observed decrease in Vmax at higher Fab5 concentrations indicates a non-competitive inhibition mechanism. (d) SEC-MALS analysis showing that Fab5 stabilizes the monomeric form of wild-type HCMV Pr. Dashed lines in panels (d) and (e) indicate expected masses of each protein species. (e) SEC-MALS analysis demonstrating that Fab5 also isolates the dimer-promoting mutant S225M HCMV Pr in a monomeric state.

We previously identified small molecule inhibitors that mimic the chemical properties of helix α5 to disrupt dimerization and decrease viral infectivity in cells, but developing high-potency leads has proven challenging^5,11,18^. We therefore sought to expand our repertoire of HHV Pr inhibitors by developing new approaches to dimer disruption. Recombinant antibodies are well suited for this task because they have been used to discover novel mechanisms of protease inactivation, including non-canonical substrate pocket binding, oligomerization disruption, and allosteric site modulation^19–22^. Antibodies that stabilize monomeric HHV Pr can also shed light on the molecular mechanisms underpinning the disorder-to-order transition to better understand how dimerization leads to catalytic activity.

Here, we describe the discovery of an antibody fragment of antigen binding (Fab) termed Fab5 that inhibits HCMV Pr by preventing dimerization through a previously unknown mechanism. High-resolution cryoelectron microscopy (cryo-EM) reveals that Fab5 binds a cryptic site distal from the active site and dimer interface, sequestering a latch loop that normally secures the C-terminal tail during the activation process. This latch loop is structurally conserved across HHV Prs, and a critical 3-residues motif is present five of eight HHV family members^11,13,14,23–26^. Mutagenesis of this motif in both HCMV Pr and KSHV Pr disrupts dimerization and neutralizes activity, revealing a conserved regulatory mechanism that can be targeted for antiviral intervention.

### Fab5 is a Non-Competitive Inhibitor that Isolates HCMV Pr Monomers

Using a phage-displayed naïve human B-cell derived Fab library, we identified five unique Fabs that bind HCMV Pr (Fig. 1b, Supplementary Fig. 2). We immobilized HCMV Pr on magnetic beads at low protein concentration to bias binders toward the monomeric species^27,28^. To monitor *in vitro* HCMV Pr activity in the presence of each Fab, we used a peptide substrate with an internally quenched fluorophore. We selected our best inhibitor, Fab5 (IC_50_ = 722 nM, Fig. 1b), for further study. Fab5 binds HCMV Pr with a K_d_ of 1.6 μM as measured by biolayer interferometry (BLI, Supplementary Fig. 3a).

We performed steady-state kinetics assays of HCMV Pr in the presence of increasing concentrations of Fab5 and fit the results to the Michaelis-Menten equation (Fig. 1c). The mechanism of Fab5 inhibition was evaluated using a Lineweaver-Burk plot (double-reciprocal plot) (Supplementary Fig. 3c); the V_max_ of the reaction decreases, and the Km modestly increases at higher Fab5 concentrations. These data are consistent with a non-competitive mode of inhibition meaning that Fab5 does not directly compete with the substrate at the active site (Supplementary Fig. 3d)^29^. This was the first evidence that Fab5 is an allosteric inhibitor of HCMV Pr.

To test whether Fab5 disrupts HCMV Pr dimerization, we determined the stoichiometry between Fab5 and HCMV Pr. Using Size-Exclusion

Chromatography/Multi-Angle Light Scattering (SEC/MALS), we measured the molar mass of the Fab5**/**HCMV Pr complexes to determine whether Fab5 binds HCMV Pr monomers or dimers. The measured mass of Fab5 alone was 47 kDa (Supplementary Fig. 3b) and HCMV Pr wild type (WT) was 32 kDa (Fig. 1d). The observed mass of the complex was 72 kDa, which is approximately the expected mass of one Fab5 bound to one HCMV Pr monomer (Fig. 1d). We repeated this experiment with HCMV Pr S225M, a mutant that promotes dimer formation (Fig. 1e)^30^. SEC/MALS of HCMV Pr S225M alone results in a single peak corresponding to a dimer (55 kDa), yet the Fab5/HCMV Pr complex sequesters HCMV Pr S225M monomers and elutes as a single 70 kDa peak. We hypothesize that the measured mass of Fab5/HCMV Pr complexes is slightly lower than expected because unbound Fab5 and HCMV Pr may co-elute with the complex^31^. Overall, these results demonstrate that Fab5 isolates HCMV Pr monomers.

### Cryo-EM Structures of Fab5/HCMV Pr Complex Reveal a Cryptic Site

To provide structural insights into how Fab5 isolates HCMV Pr monomers, we used cryo-EM to examine their interaction. We determined the structure of the Fab5/HCMV Pr complex where Fab5 and its epitope are resolved to 2.6 Å (Fig. 2a-b, Supplementary Fig. 5). Fab5 binds the latch loop, which is positioned distally from both the active site and dimer interface (Fig. 2c). HCMV Pr residues 111-117 comprise the primary epitope, with Fab5 heavy chain complementarity determining regions (CDRs) making additional contacts with the HCMV Pr core (Fig. 2d). When HCMV Pr is dimeric, the latch loop closes on the C-terminal tail and secures it against the core of the protein (Fig. 2e). Fab5 recognizes an open latch loop conformation and displaces the C-terminal tail (Fig. 2f). Importantly, Fab5 does not block the positions where helices α5 and α6 would form in a dimer, yet no EM density corresponding to these helices appears even at low thresholds (Fig. 2g). This suggests that disrupting the interaction between the C-terminal tail and the latch loop prevents the entire C-terminal region from folding into its ordered, catalytically active form.

**Fig. 2:**
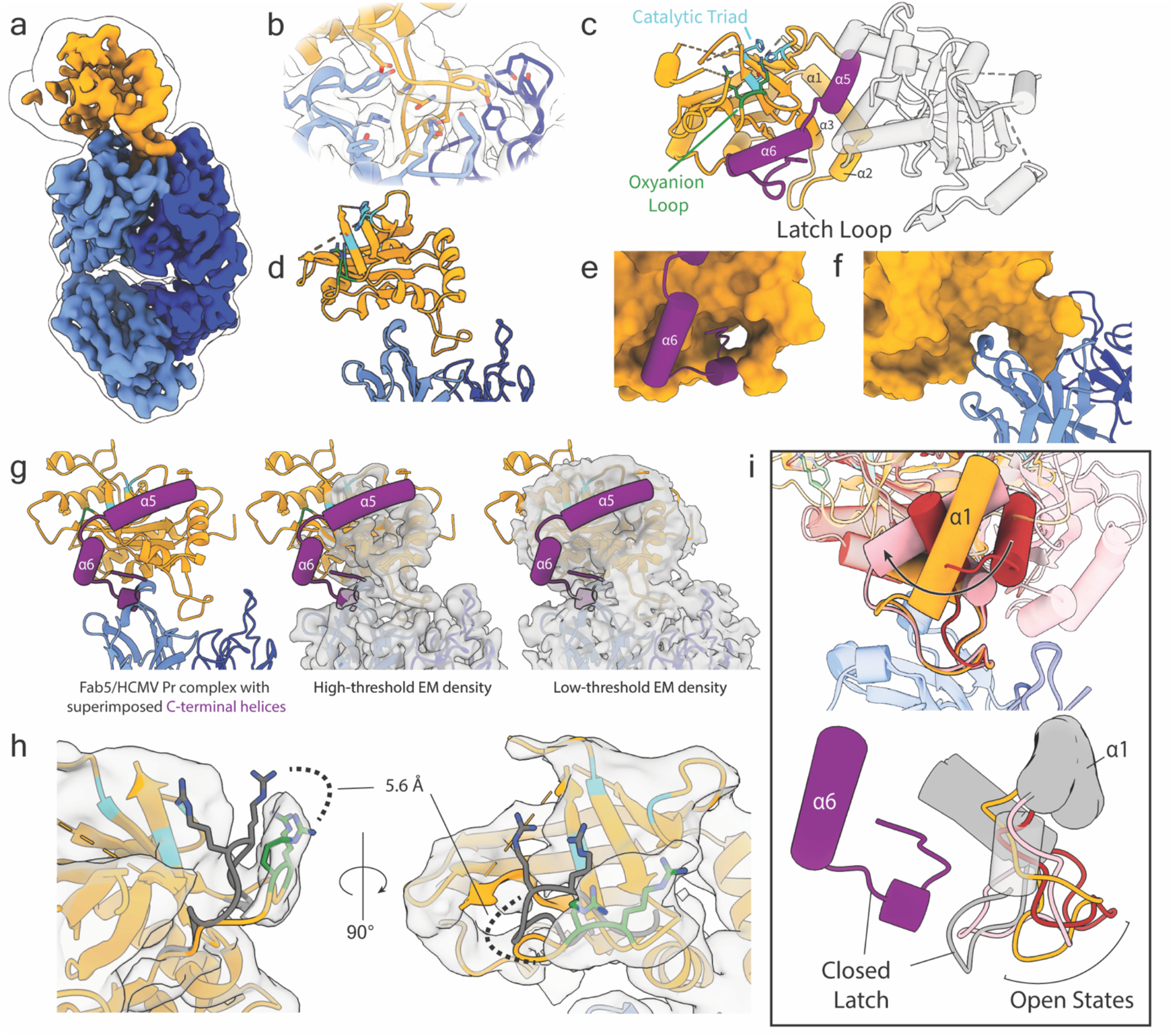
Fab5 binds the HCMV Pr latch loop. (a) Cryo-EM density map of the Fab5/HCMV Pr complex. The Fab5 light chain appears in dark blue, the heavy chain in cornflower blue, and HCMV Pr in orange. The outline contour represents EM density at a lower threshold. (b) Close-up view of the Fab5 epitope. Transparent EM density surrounds the atomic model. (c) Reference dimeric crystal structure of HCMV Pr (PDB:1CMV) with labels for the active site. (d) Atomic model of the Fab5/HCMV Pr complex highlighting the interface. (e) View of the latch loop (orange surface) in a closed conformation from dimeric HCMV Pr bound to the C-terminus in PDB:1CMV. (f) The latch loop in its open conformation when bound to Fab5. (g) HCMV Pr C-terminal region from reference dimeric crystal structure (purple tube helices) overlaid on the Fab5/HCMV Pr complex. EM density is shown at high (second panel) and low (third panel) thresholds. (h) Low-resolution EM density indicating OHL opening with respect to the dimer (gray). The distance is measured from the alpha carbon of residue R165. (i) 3D classification revealed 3 discrete poses of HCMV Pr (colored orange, red, or pink) relative to Fab5. In panel 1, structures are aligned to Fab5 and show helix α1 twisting up to 65° and tilting up to 23°. In panel 2, structures are aligned to PDB:1CMV (gray and purple). The Fab5-bound latch loops show various states of opening compared to the crystal structure.

The resolution of HCMV Pr density decreases with distance from Fab5, likely due to flexibility between the two components of the complex. Although the active site is poorly resolved, we can infer the orientation of the OHL from low-resolution density. The OHL appears to open by 5.6Å (measured from the α-carbon of R165) (Fig. 2h), resembling the OHL conformation found in inhibited KSHV Pr crystal structures where helix α6 is also absent (Supplementary Fig. 4)^18^. This confirms that proper OHL positioning requires ordering of the C-terminal region achieved through dimerization.

3D classification revealed two additional poses of HCMV Pr relative to Fab5, allowing us to observe the latch loop in various states of opening (Supplementary Fig. 5, 6). When bound to Fab5, HCMV Pr exhibits considerable range of motion, allowing for 65º of rotational twist and 23º of tilt (Fig. 2i panel 1, Movie 1). Aligning these poses to the dimeric HCMV Pr crystal structure reveals how the latch loop adopts conformations ranging from partially to fully open (Fig. 2i panel 2, Movie 2). This flexibility supports our hypothesis that the latch loop is a dynamic element that stabilizes the conformational transitions required for protease activation.

### The Latch Loop Governs HCMV Pr Dimerization and Activity

We investigated the functional role of the latch loop by performing site-directed mutagenesis of key residues and measuring HCMV Pr activity and dimerization. The interface between the latch loop and the HCMV Pr C-terminus features a salt bridge between E122 and K255, while Y253 inserts into a hydrophobic pocket formed by the latch loop with its hydroxyl group remaining solvent-exposed (Fig. 3a). Alanine substitution of any of these three latch motif residues abolishes HCMV Pr enzymatic activity in steady-state kinetics assays (Fig. 3b). We evaluated the effect of these alanine point mutations on dimerization by using SEC, which clearly separates HCMV Pr monomers and dimers into distinct peaks (Supplemental Fig. 7). We quantified the monomer-dimer equilibrium across a concentration range from 0.9 μM to 35 μM by calculating the ratio between the area under the curve of these peaks (Fig. 3c). The E122A mutant only dimerizes at the highest tested concentration (35 μM), while Y253A and K255A mutants remain exclusively monomeric throughout the entire concentration range.

**Fig. 3:**
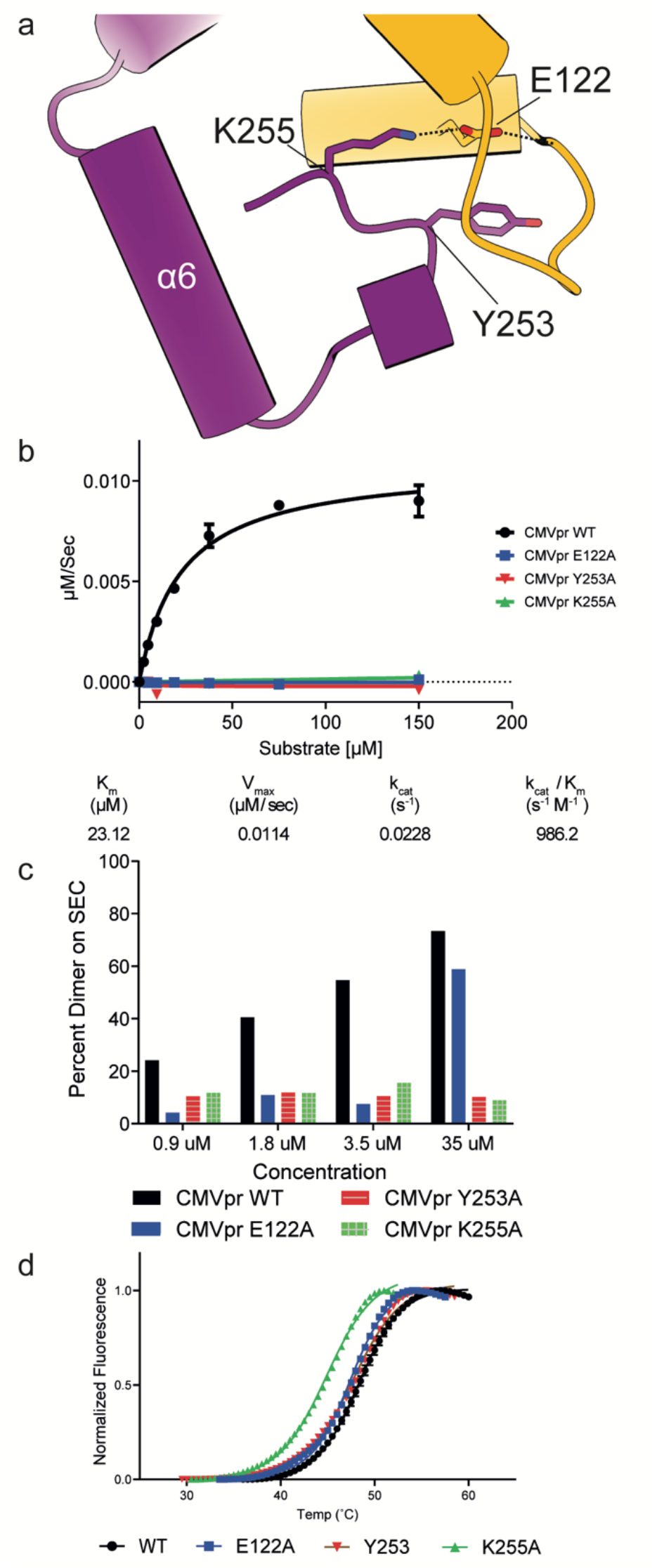
The HCMV Pr latch loop plays an important role in dimerization and activity. (a) Residues E122, Y253, and K255 form critical contacts between the latch loop and the C-terminus and comprise the latch motif. (b) Alanine substitution of any latch motif residue abolishes enzymatic activity as demonstrated by steady-state kinetics assays. (c) Concentration-dependent dimerization of wild-type HCMV Pr as measured by SEC. Percent dimerization is quantified by the ratio of integrated peak areas corresponding to monomeric and dimeric species. The E122A mutant shows dimerization only at the highest tested concentration, while Y253A and K255A mutants fail to form dimers at any concentration. (d) Thermal stability profiles of HCMV Pr mutants assessed by DSF. The K255A mutation decreases thermal stability by 4°C compared to wild-type. Other mutations induce only minor shifts in melting temperature.

To determine whether these mutations were globally destabilizing or specifically modulating dimerization equilibrium, we measured the thermal stability of the latch mutants using differential scanning fluorimetry (DSF). These experiments were performed at an HCMV Pr concentration of 4µM, where both monomeric and dimeric species of the wild-type protein would normally be present. The mutants exhibited only modest decreases in apparent melting temperature compared to wild-type HCMV Pr, with K255A showing the largest reduction (−3.4°C). Other mutations decreased thermal stability by 1°C or less (Fig. 3d). These results indicate that the latch mutations do not substantially compromise the overall protein fold or stability, suggesting their effects on protease activity and dimerization are mechanistic rather than structural. Collectively, our findings establish that the C-terminal latch plays a critical role in regulating both HCMV Pr dimerization and catalytic activity.

### Structural and Sequence Conservation of the HCMV Pr Latch Across the HHV Pr Family

The eight HHVs are classified into three subfamilies (α, β, and γ). Though HCMV Pr has a sequence similarity of only 24-38% compared with other HHV Prs, all known HHV Pr structures have high structural conservation, including the latch loop. Despite this sequence divergence, the three-residue HCMV Pr latch motif can be found in all β and γ HHV Prs (Fig. 4a). Structural alignment of HCMV Pr, KSHV Pr and Eppstein-Barr virus protease (EBV Pr) indicates that the latch motif forms the interface between the latch loop and the C-terminus across these two HHV subfamilies (Fig. 4b). We assessed the conserved role of these residues with alanine substitution of the latch motif in KSHV Pr, which results in an inactivated enzyme as it does in HCMV Pr (Fig. 4c). This functional conservation of the latch loop across HHV subfamilies reveals a fundamental mechanism regulating HHV Pr activation.

**Fig. 4:**
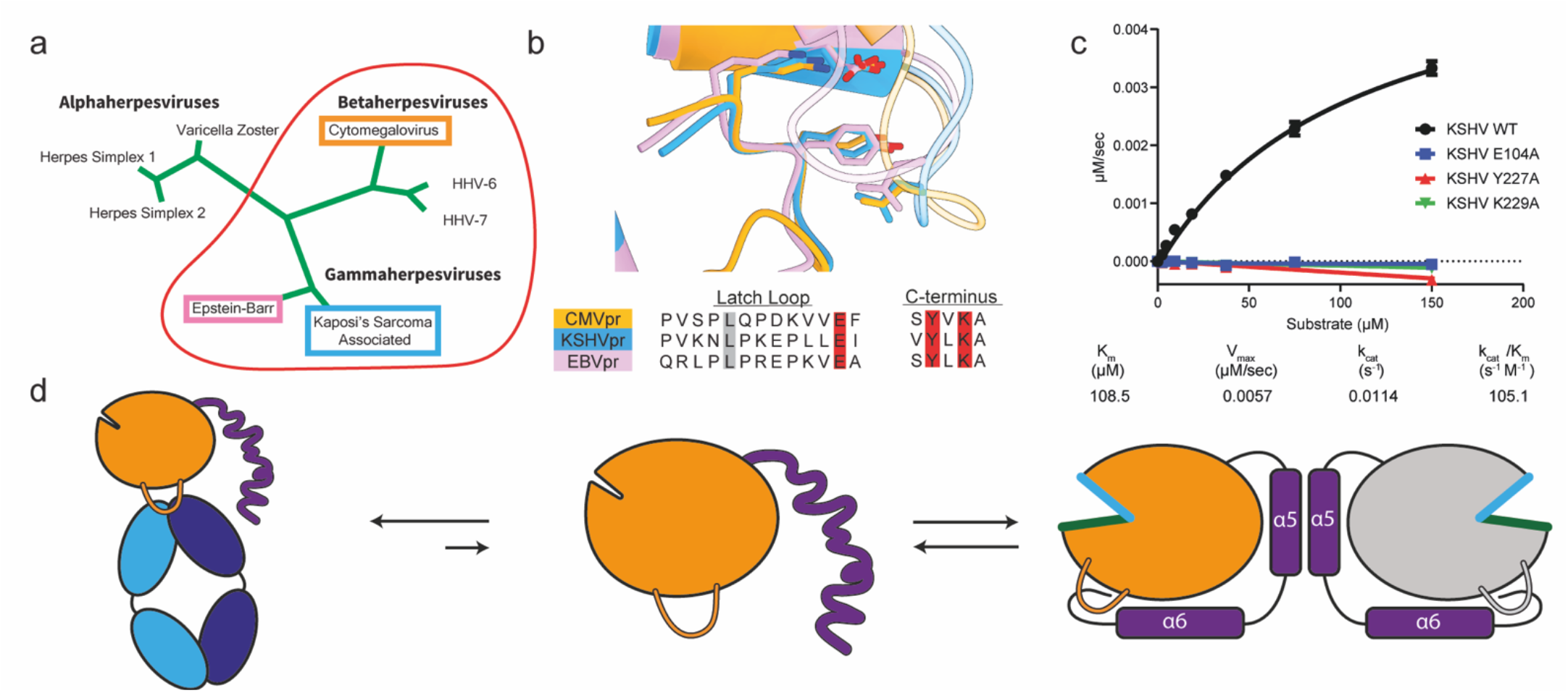
The Latch Loop is structurally and functionally conserved across the HHV Family. (a) Residues E122, Y253, and K255 form critical contacts between the latch loop and the C-terminus and comprise the latch motif. (b) Alanine substitution of any latch motif residue abolishes enzymatic activity as demonstrated by steady-state kinetics assays. (c) Concentration-dependent dimerization of wild-type HCMV Pr as measured by SEC. Percent dimerization is quantified by the ratio of integrated peak areas corresponding to monomeric and dimeric species. The E122A mutant shows dimerization only at the highest tested concentration, while Y253A and K255A mutants fail to form dimers at any concentration. (d) Thermal stability profiles of HCMV Pr mutants assessed by DSF. The K255A mutation decreases thermal stability by 4°C compared to wild-type. Other mutations induce only minor shifts in melting temperature.

## Discussion

The latch loop regulates HHV Pr dimerization with a conserved mechanism across the viral family. The equilibrium between HHV Pr monomers and dimers depends on a disorder-to-order transition of the C-terminus, with the latch loop stabilizing this structural rearrangement. Fab5 inhibits HCMV Pr activity by sequestering the latch loop and preventing it from securing the C-terminal tail (Fig. 4d). This cryptic site is only available on HCMV Pr monomers, forming the basis for the conformational selectivity of Fab5.

Structural characterization using X-ray crystallography has been a pivotal tool for discovery of compounds that bind HHV Pr and establishment of the disorder-to-order transition model^11,13,14,23–26^. However, structural information showing this disorder-to-order transition has remained elusive because the high protein concentrations required for crystallization make it difficult to study HHV Pr monomeric conformations and its many flexible loops. Using single-particle cryo-EM, we examined the Fab5/HCMV Pr complex at concentrations 100-fold lower than those required for crystallography and characterized its heterogeneity and flexibility. This work reinforces the enabling role of Fabs as conformationally selective fiducial markers^32^ and demonstrates that there should be no size barrier for Fab epitope mapping via cryo-EM^33^.

The latch loop exemplifies long-range allosteric regulation in enzyme function. By disrupting this peripheral loop, Fab5 destabilizes the active site by preventing the ordering of large structural elements. Additionally, entropy may also contribute to the Fab5 inhibitory mechanism. We recently provided comprehensive evidence linking cryo-EM density loss and B-factor changes to conformational flexibility and entropy redistribution in protein complexes^34^. Namely, rigidification of a protein interface can lead to increased flexibility in distal areas. Molecular dynamics studies of homologous KSHV and Varicella Zoster virus (VZV) proteases corroborate this model by indicating that dimerization stabilizes the dimer interface and increases mobility at the substrate binding pocket^35^. Fab5 binding may introduce flexibility in both the active site and the C-terminal region, as indicated by the respective loss of EM density. This general increase in flexibility may be an entropic barrier that prevents helices α5 and α6 from ordering even though Fab5 would sterically permit for their rearrangement.

Other allosteric serine protease inhibitors such as the anti-human Kallikrein-related peptidase Fab (anti-hKLK5) highlight how antibodies can be potent and specific by exploiting unique mechanisms of inhibition^36^. Fab5 serves as a tool compound demonstrating how the challenges of targeting the HCMV Pr active site can be circumvented via latch loop binding. Though latch loop sequences diverge among the HHV Prs, the pocket formed by the latch loop has similar properties across the viral family, suggesting that a broad-spectrum antiviral could be developed by targeting this site. Antibodies have been shown to be specific and fast-acting in the cytoplasm of human cells via electroporation, suggesting that antibody-based therapeutics targeting HHV Prs could be viable if the challenge of cellular delivery is overcome, and that feasibility is expanding due to advances in nanoparticle protein carriers and adeno-associated virus gene transfer^37–39^. While Fab5 likely lacks the potency to test this hypothesis, our structures provide the blueprint to rationally design more potent inhibitors, and Fab-phage panning against other HHV Prs would expand inhibitor repertoires even further and may allow for convergence on a cross-reactive antibody. More broadly, our findings demonstrate that the latch loop is a conserved regulatory element that can be exploited to combat HHV replication. The latch loop mechanism highlights how dynamic conformational changes can control enzyme function and underscores the importance of characterizing cryptic sites.

## Materials and Methods

### Phage display panning

A fully human naïve B-cell Fab-phage library was used for panning as previously described^40^. HCMV Pr was biotinylated using EZ-Link NHS-Chromogenic-Biotin (Pierce) and immobilized using magnetic streptavidin beads (Invitrogen) at a concentration of 100 nM. Four rounds of panning were performed with decreasing concentrations of HCMV Pr (50 nM, then 25nM) along with a negative selection using uncoated magnetic beads. Candidate Fabs were sequenced, and unique clones were expressed in BL21(DE3) *E. coli* for subsequent analysis.

### HCMV Pr constructs and mutants

All HCMV Pr constructs bear a N-terminal 6X-His tag in addition to the mutations A141V, A143V, P144A and A209V to protect against autoproteolysis. Point mutagenesis for latch motif residues was performed by designing primers with the desired point mutation in the center of the primer and predicted annealing temperatures of 68-72 ^°^C. PCR reactions were prepared by mixing 1 ng of template with 0.5 uM final of primers in ultrapure water and finally 2x Q5 master mix (NEB). PCR was performed by first holding the mixture at 95C for 3 minutes, then 28 cycles of 95C for 10 seconds, 15seconds of annealing at various temperatures, and extension at 72C for 2.5 minutes. Final extension was done for 3 minutes at 72C. Samples were then digested with DpnI (NEB) for 1 hour at 37C, then transformed into DH5-alpha cells (NEB). Cultures were grown in 5 mL Luria Broth (LB) supplemented with ampicillin (100 μg/mL final concentration) while shaking at 37^°^C overnight. DNA was isolated using a Qiagen Miniprep Kit.

### Protein Purification

HCMV Pr, KSHV Pr, and their respective latch motif mutants were expressed in Rosetta 2 BL21 *E. coli* cells (Millipore). Cells were grown in 50 mL LB supplemented with AMP (100 μg/mL) while shaking at 37^°^C overnight. The next day, 10-50 mL of culture was used to inoculate 1 L LB supplanted with ampicillin (100ug/mL) and shaken at 37^°^C to an OD_600_ of 0.6. Isopropyl β-d-1-thiogalactopyranoside (IPTG) was added (1 mM final concentration) and cultures were shaken at 16^°^C overnight. Cells were harvested, pelleted and suspended in a buffer containing 50 mM potassium phosphate at pH 8.0, 300 mM KCl, 25 mM imidazole and 5 mM 2-mercaptoethanol (BME). Cells were lysed by microfluidization and pelleted, and the supernatant was purified on a GE HealthCare LifeSciences Akta Explorer FPLC at 4^°^C. Protein was eluted over two stacked 5 mL HisTrap Nickel columns with gradient elution into a buffer containing 25 mM potassium phosphate at pH 8.0, 150 mM KCl, 300 mM imidazole and 5 mM BME. Eluate was collected and dialyzed overnight against a buffer containing 25 mM potassium phosphate at pH 8.0, 150 mM KCl, 0.1 mM EDTA and 1 mM BME. Dialyzed protein was concentrated to ∼2 mL and purified over HiLoad 26/60 Superdex 75 (GE Healthcare) into the same buffer. Protein bands were analyzed by SDS-PAGE and pure protein was collected, flash frozen and stored at −80^°^C. Protein concentrations were determined using a NanoDrop 2000c UV spectrophotometer (Thermo Scientific) using an extinction coefficient of 28,420 M^-1^ cm^-1^ for all HCMV Pr constructs. Fabs 1-5 were expressed and purified using nickel-NTA affinity columns as previously described.^28^

### Steady-state kinetics

HCMV Pr wild-type or mutant was diluted to 500 nM in buffer containing 25 mM potassium phosphate at pH 8.0, 150 mM KCl, 0.1 mM EDTA, 1 mM BME, 10% glycerol and 0.01% TWEEN-20. An internally quenched FRET substrate was used: NH_2_-Lys(MCA)-Tbg-Tbg-Asn-Ala-Ser-Ser-Arg-Leu-Lys(Dnp)-Arg-OH, where Tbg is L-*tert*-leucine, Lys(MCA) is a lysine residue with side chain linked to a 7-methoxycoumarin-4-acetic acid and Lys(Dnp) is a lysine linked to 2,4-dinitrophenyl. Substrate was prepared in 1:2 serial dilutions from 7.5 – 0.06 mM in DMSO, then 98μL protein + 2 μL substrate (150 – 0.3 μM final concentration) in a 96-well black untreated polystyrene plate. Enzyme velocity was monitored by fluorescence increase (excitation: 328 nm, emission: 393 nm) on a BioTek Synergy H4 Bioreader at 30^°^C. Mean velocity (RFU/s) during steady state was fit using BioTek Gen5 data analysis software. The mean velocity of control wells containing only buffer and substrate was averaged and subtracted from calculated enzyme velocity. A serial dilution of free MCA was prepared, ex/em at 30^°^C was determined and fluorescence (RFU) was plot in triplicate against MCA concentration (μM) then fit to a linear regression, which was used to convert initial velocity values from RFU/s to μM/s. Substrate concentration vs. velocity curves were plot in GraphPad Prism then fit using the standard Michaelis-Menten and *k*_*cat*_ equations then resulting values were used to calculate *k*_*cat*_/K_m_ with standard propagation of error. All data were collected duplicate or triplicate from technical replicates and are reported in figures as the mean, including error bars depicting standard deviation.

### IC_50_ values

HCMV Pr was diluted to 1 μM in buffer containing 25 mM potassium phosphate at pH 8.0, 150 mM KCl, 0.1 mM EDTA and 10% glycerol. Fab 1, 2, 3, 4 or 5 was diluted in 1:3 serial dilution from 8 - 0 μM in phosphate-buffered saline (PBS) then 48 μL HCMV Pr solution and 50 μL Fab solution was combined in a 96-well black untreated polystyrene plate and incubated for 20 min at room temperature (500 nM HCMV Pr final, 5% glycerol). A 7-amino-4-carbamoylmethylcoumarin (ACC) cleavable substrate was used: Ac-NH-Tbg-Tbg-N4Me_2_Asn-Ala-ACC, where Ac-NH refers to an acyl capped N-terminus, Tbg is L-*tert-*Leucine and N4Me_2_Asn is N4,N4-dimethyl asparagine. Substrate diluted in DMSO was added to the HCMV Pr + Fab solution (2 μL, 40 μM final) and enzyme velocity was monitored by fluorescence increase (excitation: 380 nm, emission: 460 nm) on a BioTek Synergy H4 Bioreader at 30^°^C. The velocity (V) of PBS-only controls was averaged and used to normalize data as percent activity:

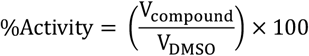

The log_10_[Fab] was plotted vs. percent activity and curves were fit with a standard four parameter log(inhibitor) vs. response with variable slope equation.

### Biolayer-interferometry

HCMV Pr was diluted into PBS then biotinylated for 1 h at room temperature using EZ-Link NHS-Chromogenic-Biotin (Pierce).^28^ Fab5 was dialyzed into buffer containing 25 mM potassium phosphate at pH 8.0, 150 mM KCl and 0.1 mM EDTA then diluted to 10,000 nM and serially diluted 2x into 5 more concentrations in the above phosphate buffer plus 1% bovine serum albumin and 0.01% TWEEN-20. Biotinylated HCMV Pr was immobilized on ForteBio streptavidin SA biosensors and the affinity of Fab5 for HCMV Pr was determined on an Octet RED384 biolayer interferometer as previously described.^28^

### Size-exclusion chromatography and Multi-angle light scattering

#### HCMV Pr + Fab 5

Equimolar amounts of Fab 5 and HCMV Pr were mixed and incubated at room temperature for 1 h. 500 µL of the mixture was injected onto a Superdex75 10/300 column and eluted over 1.2 CV.

#### HCMV Pr dimerization

WT HCMV Pr was diluted (0.9 μM, 1.8 μM, 3.5 μM or 35 μM final concentration) into buffer containing 25 mM potassium phosphate pH 8.0, 150 mM KCl, 0.1 mM EDTA, 10% glycerol and 1 mM β-mercaptoethanol then incubated at room temperature for 1 h. A Superdex 75 10/300 GL column was equilibrated at 4^°^C into the same buffer used to dilute proteins. For all protein samples, 500 μL was injected onto the column and protein eluted over 1 CV at 0.6 mL/min flow rate while monitoring Absorbance at 280nm. For each individual sample run, the absorbance at A280 was plotted flow volume (mL). The minimum absorbance value across the full spectrum was calculated in Microsoft Excel and subtracted from each absorbance value to correct the baseline to zero. The maximum absorbance was calculated for each sample, then all data were normalized to this value using the following equation, where A_x_ is the 280 nm absorbance at time x and A_max_ is the maximum value:

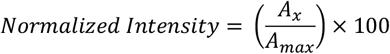

Multi-angle light scattering was performed by running the elution from an SEC run and injecting into a Wyatt Dawn Pro and Optilab combined instrument. Data were plotted in Astra and imported to GraphPad Prism where they were overlayed on normalized SEC data using a second Y-axis.

### Differential scanning fluorimetry

Protein melting temperatures were determined via differential scanning fluorimetry (DSF). DSF measurements were made using a Bio-Rad C1000 qPCR system in FRET mode. Protein (2 µM) was added to 5x SYPRO dye (Invitrogen, S6651) and plated in triplicate in a white, 96 well PCR plate in PBS. The temperature was held at 23°C for 5 min before slow ramping in 0.5°C increments every 30s. Raw data were normalized from 0 to 1 and trimmed past these minimum and maximum values before fitting in Prism. Data were fit to a Boltzmann sigmoid, and midpoints of curves are reported as melting temperature with 95% confidence interval indicated.

### Cryo-EM Sample Preparation and Data Collection

Fab5/HCMV Pr complex was purified by SEC and concentrated to 1 mg/mL. 3 microliters was pipetted onto glow-discharged gold grids coated in holey carbon film (Quantifoil, 300 mesh 1.2/1.3) or holey gold film (UltrAuFoil, 300, mesh 1.2/1.3). Grids were then blotted and plunge-frozen using a Vitrobot Mark IV equipped with Whatman type 4 blotting paper with a blotting time of 4 s and blotting force of −2, at 4 °C and 100% humidity. Data was collected at the Janelia Research Campus in Ashburn, VA on Krios 3. This 300 keV microscope is equipped with a Thermo Scientific Falcon 4i camera, Selectris X energy filter, and cold Field emission gun with 6 eV slit width during acquisition. A nominal magnification of 165,000x was used for a physical pixel size of 0.743 (0.371 in super resolution) with a total dose of 50 e-/A ° 2. Automated data collection was performed using SerialEM to collect movies with a defocus range between 0.8-1.8 Å.

### Image Processing

Dose-fractionated images stacks were motion corrected and Fourier-cropped by 2 using both cryoSPARC and MotionCor2^41^. CTF estimation was performed using cryoSPARC, followed by micrograph curation, blob-based particle picking, 2D classification, and ab initio modeling. The ab initio model was used to generate templates for further particle picking and curation. Iterative rounds of multi-class ab initio modeling and heterogeneous refinement were used to generate particle stacks where HCMV Pr could be resolved in the complex. Particles were then sent to Relion using the PyEM code suite^42^ for 3D Classification. Particles were then reimported to cryoSPARC for further 3D Classification, and 3D refinement.

### Model Building and Refinement

A model of Fab5 generated by Alphafold 3 was docked into cryo-EM density and refined using ISOLDE and PHENIX^43,44^. To identify the Fab5 epitope, we generated each flexible loop of HCMV Pr as a peptide in Coot^45^ and began modeling it into the non-Fab5 cryo-EM density *de novo*. We identified that the loop from P111-P117 fit this density well and then appended the rest of an HCMV Pr monomer onto it (PDB:1CMV) and built the missing loops in Coot. The complete model was further refined using real-space refinement in PHENIX and ISOLDE. Final collection, refinement and validation statistics are reported in supplementary table 1.

## Acknowledgements

We thank D. Asarnow, T. DeTomasi, K. Choi, and W. Choi for reading the manuscript and providing feedback. This work is supported by the National Institutes of Health (U54AI170792 to C.S.C and R35GM140847 to Y.C.). We also thank D. Bulkley, G. Gilbert, L. Want (UCSF Electron Microscopy Core) and R. Yan (Janelia Cryo-EM Facility) for their advice and assistance with cryo-EM data acquisition. Instruments at the UCSF Cryo-EM facility are partially supported by grants from the NIH (S10OD020054, S10OD021741 and S10OD026881) and Howard Hughes Medical Institute. C.S.C. received support from the UCSF Innovation Ventures (InVent) program, the UCSF Catalyst program and a generous donation from Nadav Ben-Efraim. Y.C. is an Investigator of the Howard Huges Medical institute.

## Author contributions

K.R.H. and M.F.B. discovered Fab5 through phage-displayed panning. M.Z., K.R.H., M.F.B., and J.N. performed biochemical characterization of Fab5 and HHV Pr variants. P.J.R and M.Z. performed differential scanning fluorimetry. M.Z. carried out the structural component of this study. All authors participated in data analysis and evaluation, and all authors contributed to manuscript preparation. C.S.C. and Y.C. supervised the project and provided advice, guidance, and support throughout.

## Data availability

Composite maps of the Fab5/HCMV Pr complex have been deposited into the Electron Microscopy Data Bank (EMDB) (Class 1: EMD-72658, class 2: EMD-72659, class 3: EMD-72660). Corresponding atomic models were deposited in the Protein Data Bank (PDB) (Class 1: 9Y7L, class 2: 9Y7M, class 3: 9Y7N).

## Supplementary Figures

**Supplementary Fig. 1:**
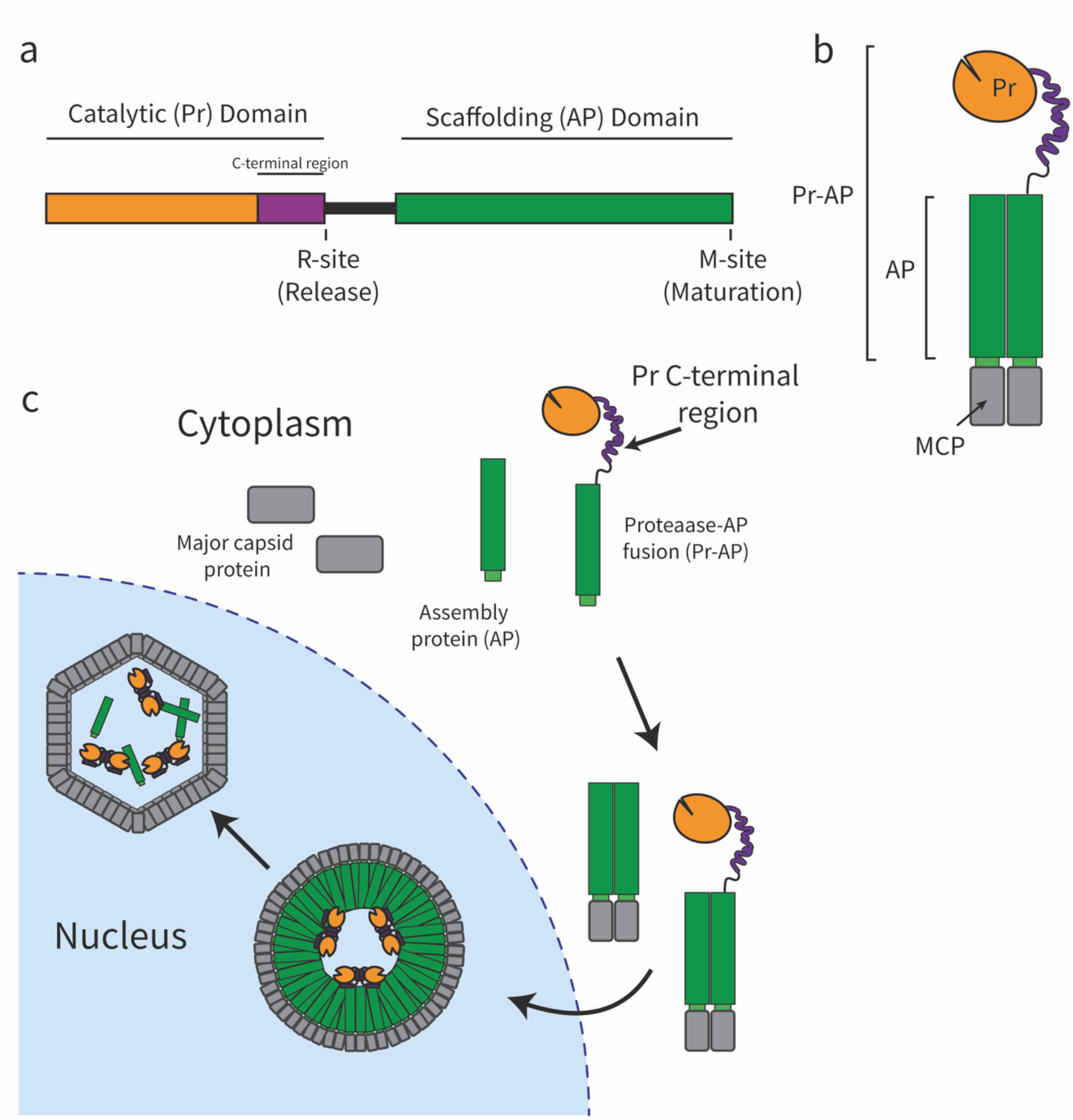
HCMV Pr gene and function. (a-b) The HCMV UL80 gene. Primary components are the catalytic domain, a linker region, and the assembly protein (AP) Scaffolding domain. Proteins can either be expressed as Pr-AP or AP due to multiple open reading frames. (c) Pr-AP associates with the major capsid protein (MCP) in the cytosol. Catalytically competent Pr frees itself from AP via the R-site, and the scaffolding is cleared away via the M-site. The wild-type Pr domain contains multiple autoproteolytic sites, which is why our expression construct contains mutations (A141V, A143V, P144A and A209V) to limit autoproteolysis.

**Supplementary Fig. 2:**
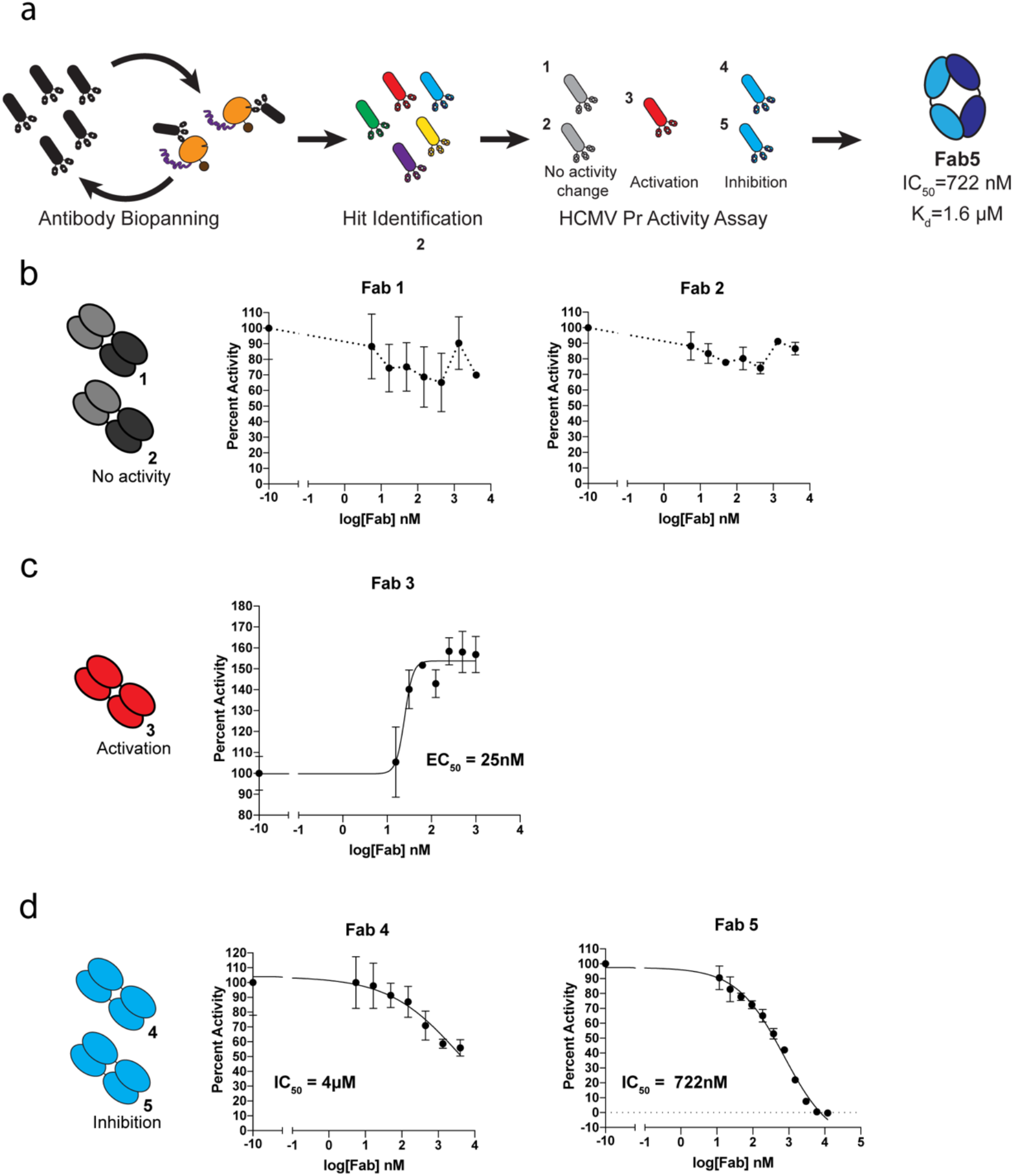
Inhibition assay with the five unique clones from HCMV Pr phage-displayed panning. (a) a schematic of the panning campaign against HCMV Pr. Five hits were expressed and purified then tested for inhibition of HCMV Pr activity. (b) Fabs 1 and 2 do not affect HCMV Pr activity. (c) Fab3 has an activating effect on HCMV Pr activity. (d) Fabs 4 and 5 inhibit HCMV Pr. We chose Fab5 for deeper characterization. We did attempt to move forward with further characterization of Fab3 as well, but we were not able to purify a stable complex on SEC or SEC/MALS.

**Supplementary Fig. 3:**
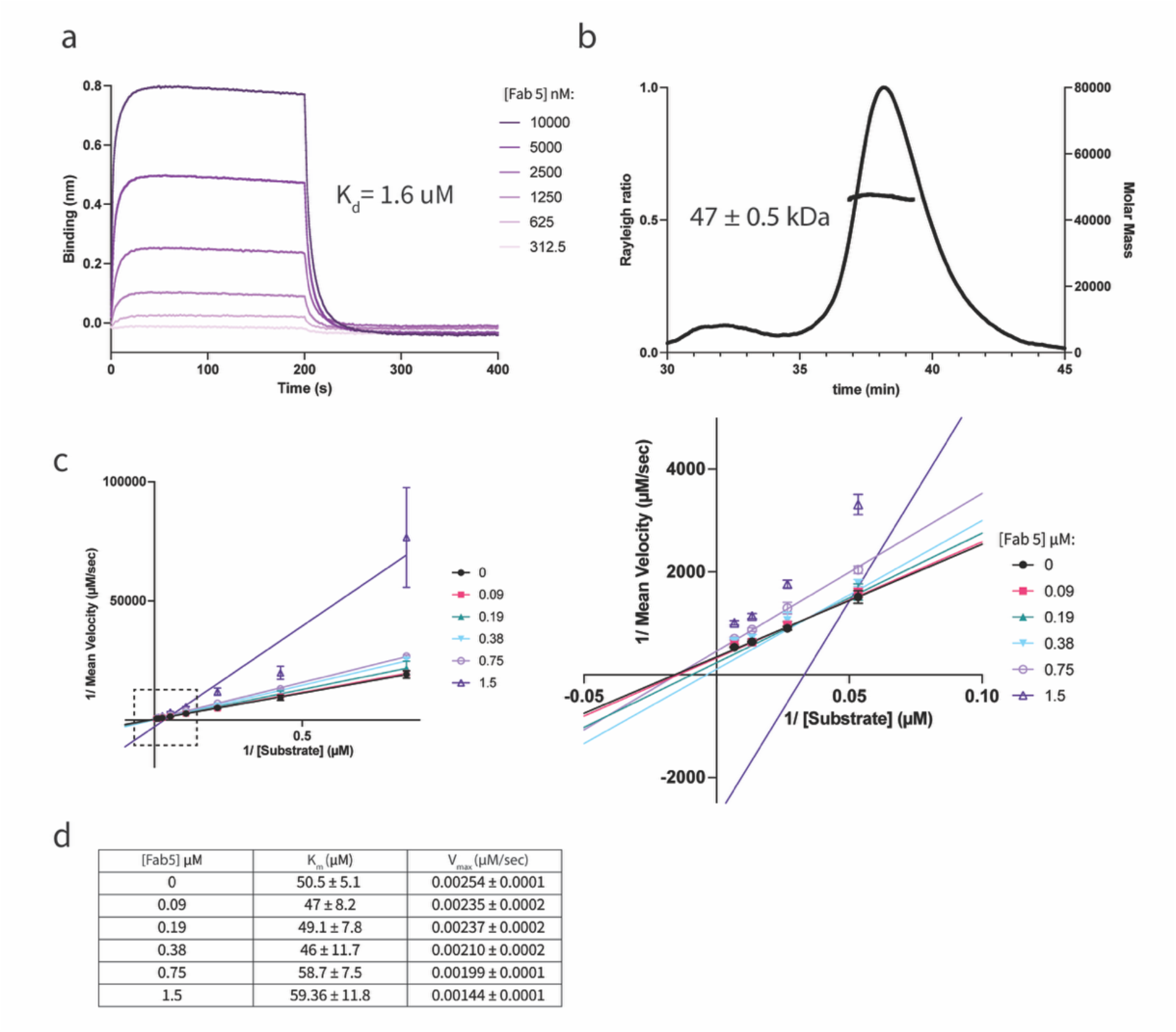
Fab5 BLI, SEC/MALS, and Steady-State Kinetics Assays. (a) Biolayer interferometry sensogram of Fab5 binding to immobilized HCMV Pr. The dissociation constant (K_d_) was determined by fitting the steady-state response to a 1:1 equilibrium binding model. (b) SEC/MALS experiment with Fab5 alone using an S200 10/300 SEC column. (c) Double-reciprocal plot of steady-state kinetics assay measuring HCMV Pr activity performed with Fab5 at various concentrations. Full data is plotted in panel 1, and a magnified view of the dotted line-boxed region is plotted in panel 2. (d) The K_m_ and V_max_ of HCMV Pr steady-state kinetics are provided in a table for each concentration of Fab5.

**Supplementary Fig. 4:**
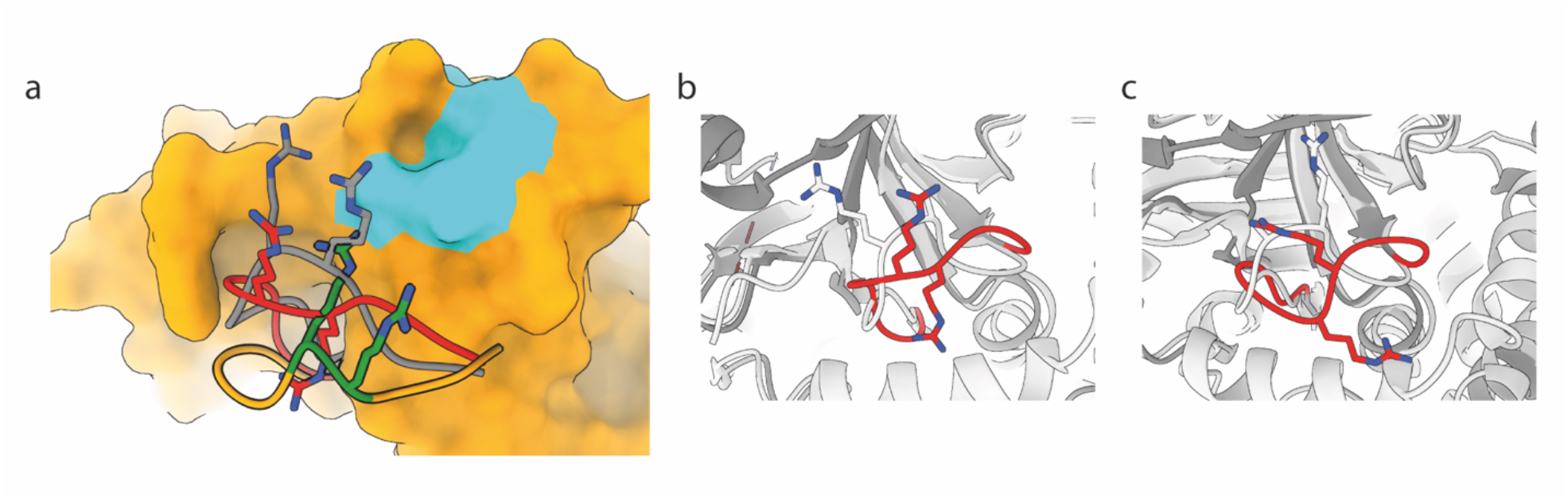
Structural comparison of the OHL. (a) HCMV Pr (orange surface) with active site labeled cyan. The three loops in the foreground are OHL loops from an HCMV Pr dimer (gray, PDB:1CMV), inhibited KSHV Pr (red, PDB:3NJQ), and the Fab5-HCMV Pr complex (orange and forest green). (b-c) The biological assembly of inhibited KSHV Pr in the crystal structure PDB:3NJQ is dimeric despite the deletion of the C-terminal region. The inhibitor, which mimics helix α5 and displaces it, forms a crystal contact between 2 monomers. The OHL assumes different conformations in chains A and B (red), however both OHL conformations are more open compared to the wild-type structure (light gray, PDB:1FL1) and the highlighted arginines point away from the catalytic triad.

**Supplementary Fig. 5:**
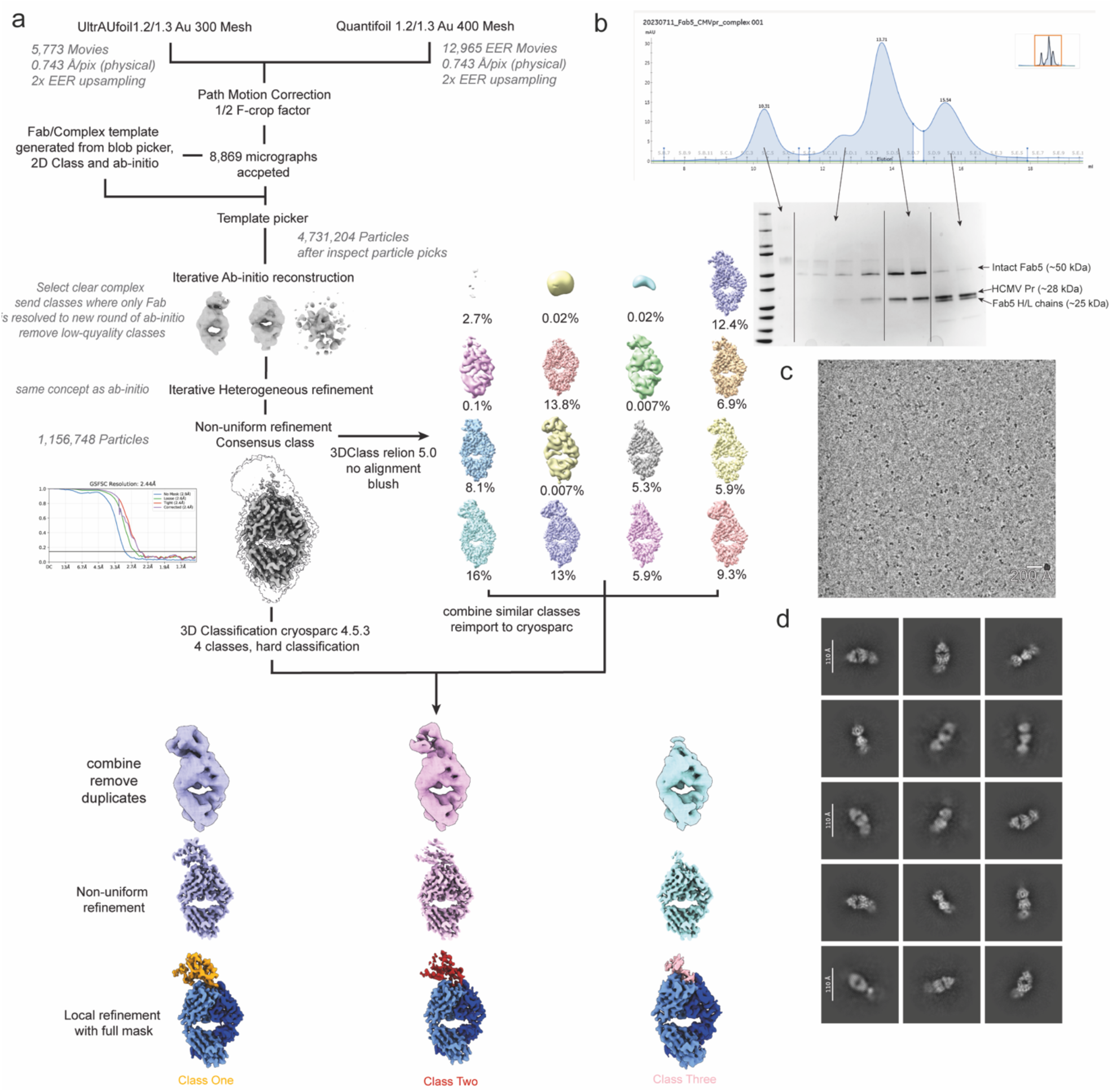
Cryo-EM Data Processing Workflow. (a) Flow chart of EM processing workflow. Final volumes for classes One, Two, and Three were solved using 3D classification, non-uniform refinement, and local refinement with a full mask. All volumes were used iteratively for model building. (b) size-exclusion chromatography of Fab5/HCMV Pr complex preparation. Only the largest peak was used for cryo-EM sample. (c) Representative micrograph of Fab5/HCMV Pr complex. (d) Representative 2D classes of Fab5/HCMV Pr complex.

**Supplementary Fig. 6:**
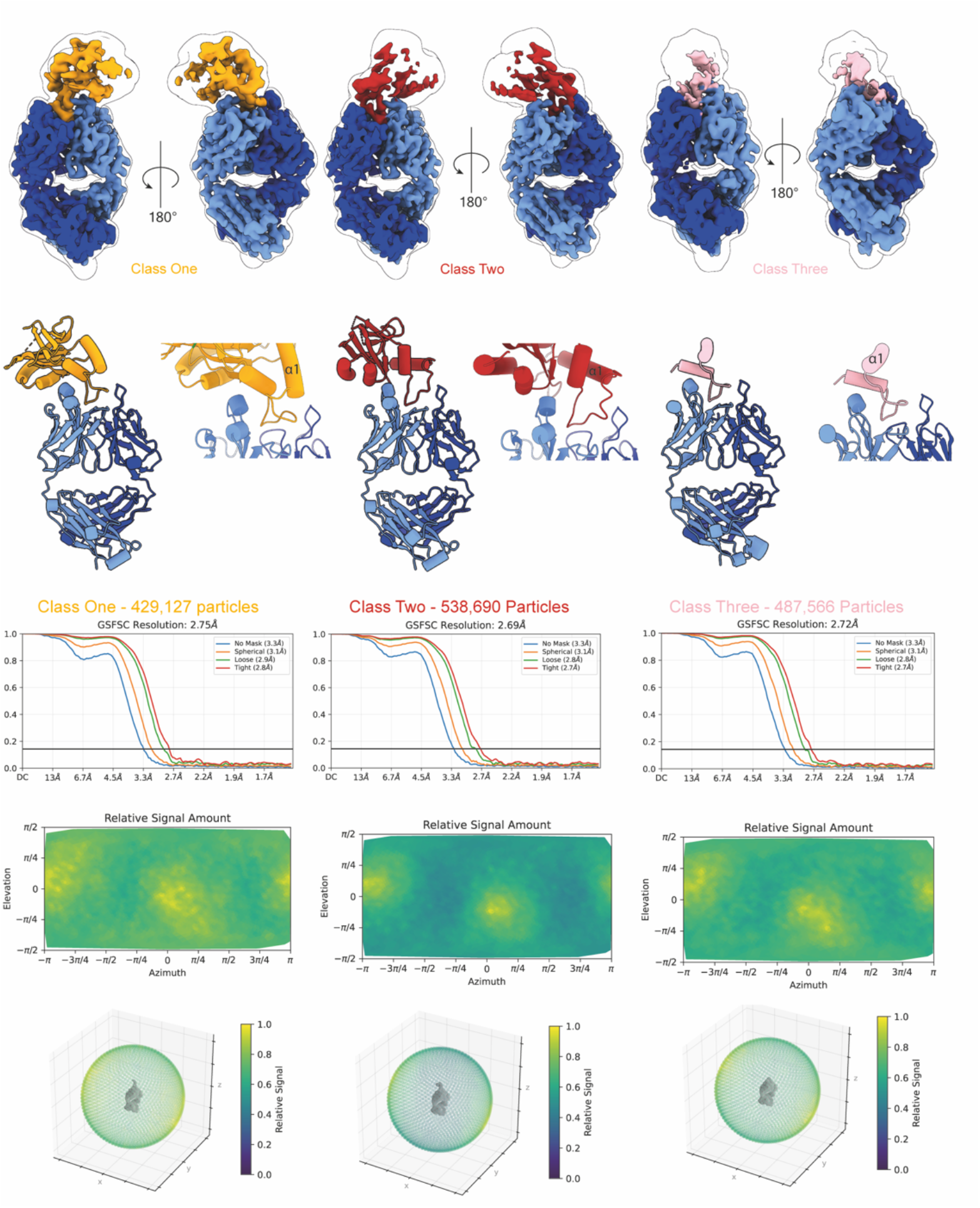
Cryo-EM Map Quality and Models. For each class, two views of the map are provided with an outline of the density at a low threshold. The descending panels show the model with an inset zoom of the epitope region, then a Fourier shell correlation (FSC) curve, then orientation diagnostics where relative signal is shown as a function of viewing direction.

**Supplementary Fig. 7:**
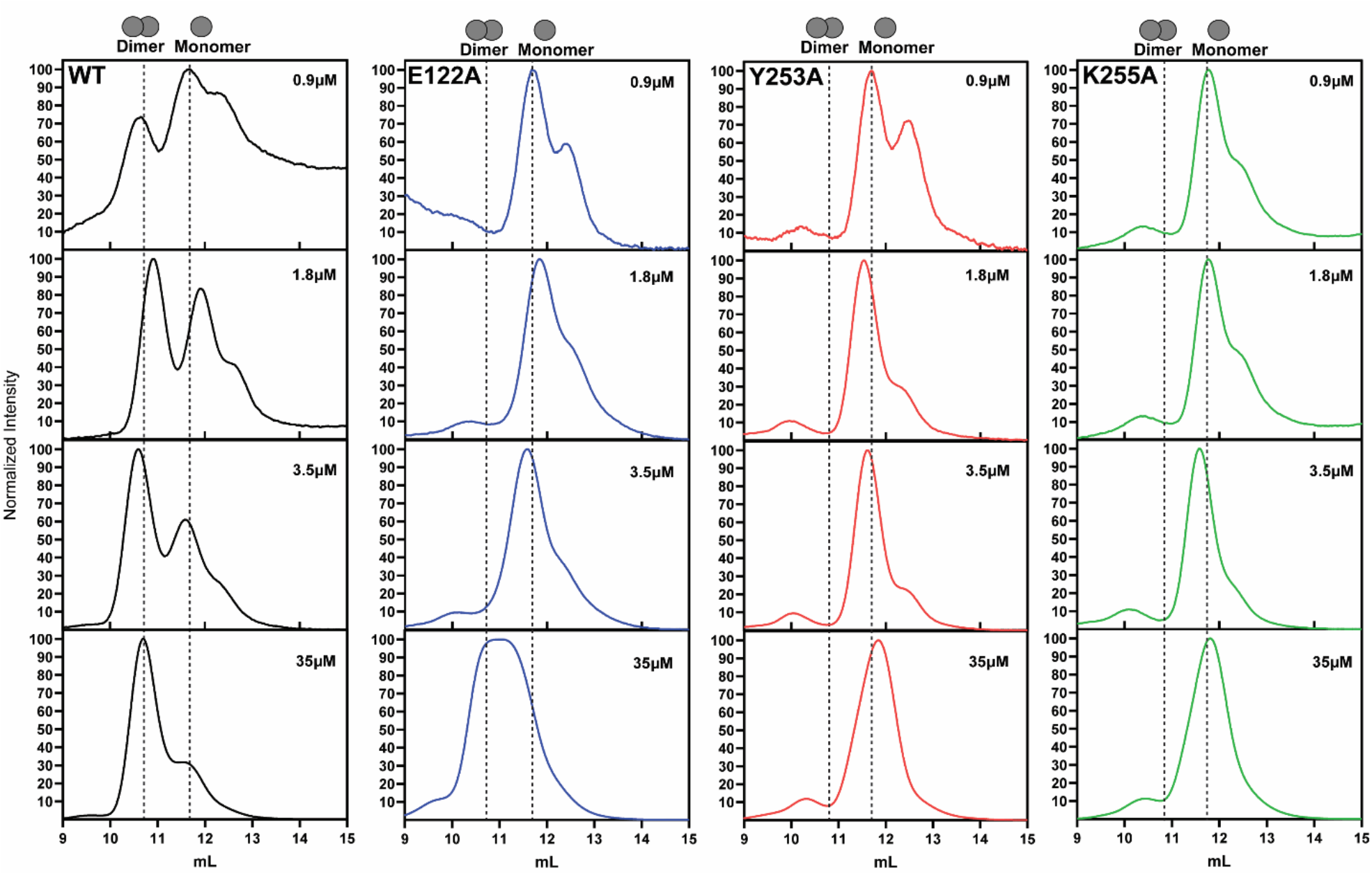
All SEC chromatograms from dimerization experiments. Peak areas were integrated, and a ratio was taken to make the percent dimer graph in Fig. 4. Results are normalized for intensity, and the x-axis has column volume. The left column is wild-type HCMV Pr, and concentration-dependent dimerization can be observed. All mutant constructs remain as monomers except for E122A, which can dimerize at the highest concentration.

**Table 1.** Cryo-EM data collection, refinement, and validation statistics.

|  | Fab5-HCMV Pr Class 1 | Fab5-HCMV Pr Class 2 | Fab5-HCMV Pr Class 3 |
| --- | --- | --- | --- |
| **Data Collection and Processing** |  |  |  |
| Magnification | EFTEM 165,000 | EFTEM 165,000 | EFTEM 165,000 |
| Voltage (kV) | 300 | 300 | 300 |
| Electron exposure (e<sup>-</sup>/Å<sup>2</sup>) | 50 | 50 | 50 |
| Defocus range (μm) | -0.8-2.0 | -0.8-2.0 | -0.8-2.0 |
| Pixel size (Å) | 0.743 | 0.743 | 0.743 |
| Symmetry imposed | C1 | C1 | C1 |
| Initial particle number | 2,795,045 | 2,795,045 | 2,795,045 |
| Final particle images | 429,127 | 538,690 | 487,966 |
| Map resolution (Å) | 2.72 | 2.57 | 2.69 |
| FSC threshold | 0.143 | 0.143 | 0.143 |
| **Refinement** |  |  |  |
| Initial model used | Ab initio | Ab initio | Ab initio |
| Model resolution (Å) | 3.3 | 3.2 |  |
| FSC threshold | 0.5 | 0.5 | 0.5 |
| Map sharpening B-factor | -117.0 | -106.1 | -114.5 |
| Model Composition |  |  |  |
| Non-hydrogen atoms | 8834 | 8772 | 7266 |
| Protein residues | 586 | 582 | 482 |
| Ligands | 0 | 0 | 0 |
| B factors (Å) |  |  |  |
| Protein (min/max/mean) | 28.40/187.37/104.02 | 75.35/336.87/136.93 | 28.40/187.37/108.92 |
| R.M.S. Deviations |  |  |  |
| Bond lengths (Å) | 0.012 | 0.012 | 0.012 |
| Bond angles (°) | 1.911 | 1.917 | 1.917 |
| Validation |  |  |  |
| MolProbity score | 0.73 | 0.75 | 0.69 |
| Clashscore | 0.11 | 0 | 0 |
| Rotamer outliers (%) | 0.40 | 0.61 | 0.45 |
| Ramachandran plot |  |  |  |
| Favored (%) | 96.87 | 96.33 | 93.70 |
| Allowed (%) | 2.96 | 3.15 | 5.25 |
| Outliers (%) | 0.17 | 0.52 | 1.05 |
| Model vs. Data |  |  |  |
| CC (mask) | 0.72 | 0.77 | 0.76 |
| CC (box) | 0.75 | 0.74 | 0.74 |
| CC (peaks) | 0.64 | 0.64 | 0.66 |
| CC (volume) | 0.73 | 0.77 | 0.77 |

